# Novel pelagiphages prevail in the ocean

**DOI:** 10.1101/2019.12.14.876532

**Authors:** Zefeng Zhang, Fang Qin, Feng Chen, Xiao Chu, Haiwei Luo, Rui Zhang, Sen Du, Zhen Tian, Yanlin Zhao

**Affiliations:** Fujian Provincial Key Laboratory of Agroecological Processing and Safety Monitoring, College of Life Sciences, Fujian Agriculture and Forestry University, Fuzhou, Fujian, China; Instituteof Marine and Environmental Technology, University of Maryland Center 10 for Environmental Science, Baltimore, MD, USA; Simon F. S. Li Marine Science Laboratory, School of Life Sciences and State Key Laboratory of Agrobiotechnology, The Chinese University of Hong Kong, Shatin, Hong Kong SAR, China; State Key Laboratory of Marine Environmental Science, College of Ocean and Earth Sciences, Institute of Marine Microbes and Ecospheres, Xiamen University, Xiamen, Fujian, China

**Author notes:** These authors contributed equally: Z.Z. and F.Q. contributed equally to this work.

**Keywords:** SAR11, pelagiphages, novel phage groups, Viromics

## Abstract

Viruses play a key role in biogeochemical cycling and host mortality, metabolism, physiology and evolution in the ocean. Viruses that infect the globally abundant marine SAR11 bacteria (pelagiphages) were reported to be an important component of the marine viral community. In this study, ten pelagiphages that infect three different *Pelagibacter* strains were isolated from various geographical locations and were genomically characterized. All ten pelagiphages are novel, representing four new lineages of the *Podoviridae* family. Although they share limited homology with cultured phage isolates, they are all closely related to some environmental viral fragments. Two HTVC023P-type pelagiphages are shown to be related to the abundant VC_6 and VC_8 viral populations of the Global Oceans Viromes (GOV) datasets. Interestingly, HTVC103P-type pelagiphages contain a structural module similar to that in SAR116 phage HMO-2011. Three HTVC111P-type pelagiphages and HTVC106P are also novel and related to GOV VC_41 and VC_67 viral populations, respectively. Remarkably, these pelagiphage represented phage groups are all globally distributed and predominant. Half of the top ten most abundant known marine phage groups are represented by pelagiphages. The HTVC023P-type group is the most abundant known viral group, exceeding the abundance of HTVC010P-type and HMO-2011-type groups. Furthermore, the HTVC023P-type group is also abundant throughout the water column. Altogether, this study has greatly broadened our understanding of pelagiphages regarding their genetic diversity, phage-host interactions and the distribution pattern. Availability of these newly isolated pelagiphages and their genome sequences will allow us to further explore their phage-host interactions and ecological strategies.

## Introduction

As the most abundant biological entities in the ocean, viruses play critical roles in impacting marine biogeochemical cycling and shaping the microbial community structure and function (1–3). They also harbor enormous genetic diversity and diverse metabolic potentials (4–7). Despite the fundamental importance of marine viruses, we just began to understand the diversity of marine viral communities. In the most recent decade, culture-independent metagenomic surveys (4–8), metagenomics fosmids (9,10) and single-cell genomics (SCGs) (11–13) have been used to explore the marine viral genetic and functional diversity and obtain novel viral genomic fragments from uncultivated viruses. For example, 15,280 viral populations belonging to 867 genus-level viral clusters were identified from the analysis of the Global Oceans Viromes (GOV) (5), 488,130 viral populations were recently identified from GOV 2.0 datasets (6), and more than 1,000 viral genomic fragments were retrieved from a single fosmid library from the Mediterranean deep chlorophyll maximum (MedDCM) (9). These studies unveiled enormous diversity of viruses in the ocean. In contrast, culture-dependent viral isolation has a limited contribution to reveal the marine viral diversity due to the fact that many bacterial groups are not easy to be cultivated in the laboratory. The number of cultured viruses from the ocean is far less compared to the number of omic-assembled viruses. The culture-independent studies have unveiled enormous diversity of viruses in the ocean, and at the same time, raised challenges on finding their potential hosts and unravelling their ecological and biological roles. Considerable efforts have been made to predict the hosts of viral sequences and to predict potential phage-host interactions (5, 9, 14). Despite these efforts, hosts of most viral clusters identified from the GOV still remain unknown and the majority of the viral clusters identified from the GOV lack any cultivated representative (5).

Although current isolated phages are insufficient for elucidating the natural viral diversity in the ocean, the isolation and genomic analysis of some important marine phages, such as cyanophages, SAR116 phage (15) and SAR11 phages (referred as pelagiphages) (16), have greatly facilitated the interpretation of marine virome datasets. The discovery of SAR11 phages (16) and a SAR116 phage (15) advocates for the importance of viral isolation. It was estimated that the isolation of pelagiphages and SAR116 phage HMO-2011 increased the number of known reads of in viral metagenomes by 30% (17). Therefore, isolation and sequencing more phages infecting ecologically important bacterial hosts is of urgency in the area of marine viral ecology. In addition, having phage isolates in culture has the advantages of obtaining full genome sequences, gaining phage biological information, establishing cultivated virus-host model systems for a better understanding the phage infection process and ecological functions in marine ecosystems.

The order *Pelagibacterales* (SAR11) within the *Alphaproteobacteria*, is ubiquitous in the marine environments, accounting for approximately one-third of the oceanic prokaryotic cells (18–20), making the SAR11 clade the largest population of closely related heterotrophic bacteria on Earth. SAR11 bacteria are typical marine “difficult–to-culture” oligotrophic bacteria, exhibiting a slow growth rate and requiring unusual culturing condition (18, 21). Due to the difficulty in SAR11 bacteria culturing and pelagiphage isolation, the genomic and ecological study of pelagiphages has just begun to be addressed by a few studies. Pelagiphages are among the most abundant marine known phage groups, and they influence the population dynamics and evolution of SAR11 (16). Given the ecological importance of SAR11, pelagiphage has received much research attention since the four pelagiphage isolates were reported in 2013. Currently, 15 pelagiphages belonging to three distinct phage groups have been reported, and they possess diverse genetic contents and novel life strategies (16, 22). Efforts are still needed to further explore the diversity of pelagiphages in order to better understand their genomic evolution and phage-host interactions.

In this study, we isolated and sequenced ten new pelagiphages from diverse marine environments. High levels of genetic diversity were revealed across these pelagiphage genomes. We showed that these pelagiphages belong to four novel viral groups and are closely related to many environmental viral fragments. Finally, metagenomic recruitment analyses reveal the dominance of certain pelagiphage represented phage groups in the upper ocean as well as in the deep ocean.

## Results and discussion

### Morphology and general features of newly isolated pelagiphages

In this study, a culture-dependent approach was employed to further explore the genetic diversity of pelagiphages. Ten pelagiphages infecting three SAR11 strains (98.9-99.4% 16S rRNA gene sequence identity) that belong to the SAR11-Ia subclade were isolated from diverse marine environments (Table 1). All pelagiphages belong to the *Podoviridae* family with capsid length ranging from 55 to 69 nm (Fig. 1*A*). It is noteworthy that among 15 previously isolated pelagiphages (16, 22), 14 are podoviruses, and only one is myovirus. No siphovirus infecting SAR11 has been reported yet. At present, all reported pelagiphages infect closely related SAR11 Ia isolates. More novel phage groups are expected to be discovered when diverse SAR11 strains are used for phage isolation.

**Table 1.** General features of pelagiphages analyzed in this study.

| Phage | Host | Source watera | Depth (m) | Latitude | Longitude | Sampling date | Phage group | Capsid size (mean±s.d., nm) | Genome size (bp) | G+C % | Number of ORFs | Accession number |
| --- | --- | --- | --- | --- | --- | --- | --- | --- | --- | --- | --- | --- |
| HTVC023P | HTCC1062 | South China Sea SEATS | 500 | S18°00' | E116°00' | Nov-2016 | HTVC023P-type | 68±2 | 60878 | 35.0 | 86 | MN698239 |
| HTVC027P | HTCC1062 | South Pole K1 | 150 | N11°39' | E78°59' | Sep-2016 | HTVC023P-type | 69±1 | 57595 | 34.8 | 86 | MN698241 |
| HTVC103P | HTCC7211 | South China Sea SEATS | 5 | S18°00' | E116°00' | Aug-2014 | HTVC103P-type | 68±1 | 54103 | 31.0 | 86 | MN698242 |
| HTVC104P | HTCC7211 | India Ocean 105 | 75 | S4°0' | E95°28' | Mar-2015 | HTVC103P-type | 67±2 | 54359 | 30.9 | 89 | MN698243 |
| HTVC115P | HTCC7211 | India Ocean 105 | 500 | S4°0' | E95°28' | Mar-2015 | HTVC103P-type | 69±3 | 54819 | 33.2 | 81 | MN698247 |
| HTVC111P | HTCC7211 | Mediterranean Sea | Surface | N43°42' | W7°17' | Aug-2016 | HTVC111P-type | 55±2 | 31577 | 30.5 | 64 | MN698245 |
| HTVC112P | HTCC7211 | South Pole K1 | 150 | S78°59' | E11°40' | Sep-2016 | HTVC111P-type | Nd | 32478 | 30.4 | 58 | MN698246 |
| HTVC026P | HTCC1062 | Mediterranean Sea | Surface | N43°42' | W7°17' | Aug-2016 | HTVC111P-type | Nd | 32480 | 31.3 | 66 | MN698240 |
| HTVC202P | FZCC0015 | Pingtang coast, Taiwan straight | Surface | N25°26' | E119°47' | May-2017 | HTVC111P-type | Nd | 32226 | 31.3 | 61 | MN698248 |
| HTVC106P | HTCC7211 | India Ocean 113 station | 500 | N0°0' | E92°59' | Mar-2015 | HTVC106P-type | 56±1 | 36945 | 32.1 | 70 | MN698244 |

ND : Not Determined.

**Fig. 1.**
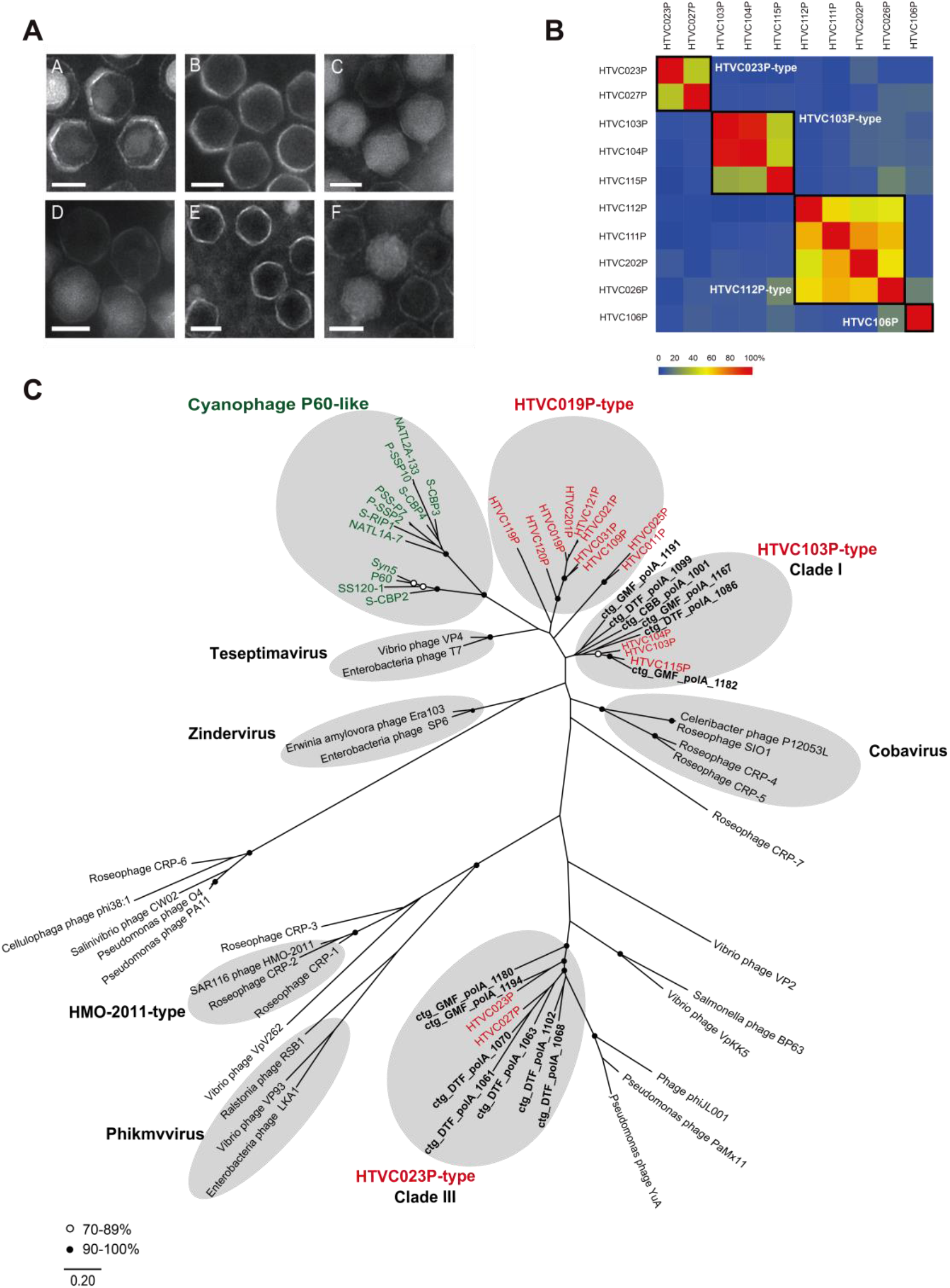
(*A*) Transmission electron microscope images of selected pelagiphages from each group. A, HTVC023P; B, HTVC027P; C, HTVC103P; D, HTVC104P; E, HTVC111P; F, HTVC106P. (Scale bars: 50 nm). (*B*) Heatmap presentation of shared genes of newly isolated pelagiphages. Phages in the same group are boxed. (*C*) Unrooted maximum-likelihood phylogenetic tree of phage DNA polymerases constructed with conserved polymerase domains. Novel DNA polymerases identified from a previous study are in bold (28). The scale bar represents the amino acid substitutions per site.

All newly isolated pelagiphage genomes were assembled into a circular contig, indicating their genome completeness. The general features of these pelagiphages are shown in Table 1. The genome sizes of these pelagiphages vary from 32.4 to 60.8 kb with a G+C content of 30.4% to 35.0%, which is close to the G+C content of their hosts (29.0 to 29.7%) and other previously reported pelagiphages (29.7 to 35.5%) (16, 22). No tRNA sequences were identified in all 10 pelagiphage genomes.

### Marine pelagiphages possess genetic diversity and novelty

Overall, the 10 pelagiphages reported here share limited sequence homology with other cultured phage isolates. Comparative genomics analysis categorized these pelagiphages into four distinct phage groups (at the genus-level approximately) (Figs. *1B* and 2). To date, seven distinct phage groups were identified from pelagiphage isolates, six of which belong to the *Podoviridae* family, suggesting that podoviruses exert primary top-down control on the abundance and dynamics of SAR11 population. The prevalence and dominance of Pelagibacter podoviruses in the ocean is supported by the dominance of their viromic matches (see later Viromic fragment recruitment analyses). Integrase genes and other genes related to the lysogenic life cycle were not identified in all 10 pelagiphage genomes, indicating that these pelagiphages all infect SAR11 bacteria using the lytic infection strategy.

### Match to the most abundant GOV viral clusters

HTVC023P and HTVC027P are closely related, belonging to a novel HTVC023P-type phage group (Fig. 2*A*). Thirty-seven ORFs are shared between HTVC023P and HTVC027P (with 31 to 93% amino acid identity) and they have a conserved overall genome arrangement with few gene rearrangements (Fig. 2*A*). HTVC023P and HTVC027P exhibit no significant genomic synteny with other known phage isolates, thus possessing genomic and evolutionary novelty. Approximately half of the predicted ORFs from both HTVC023P and HTVC027P genomes show homology to genes from other types of phage and bacterial genomes. The remaining ORFs have no homologs in the NCBI-RefSeq database and some only hit environmental sequences. Of the predicted ORFs, only 17% were assigned to putative biological functions based on the sequence homology analysis.

**Fig. 2.**
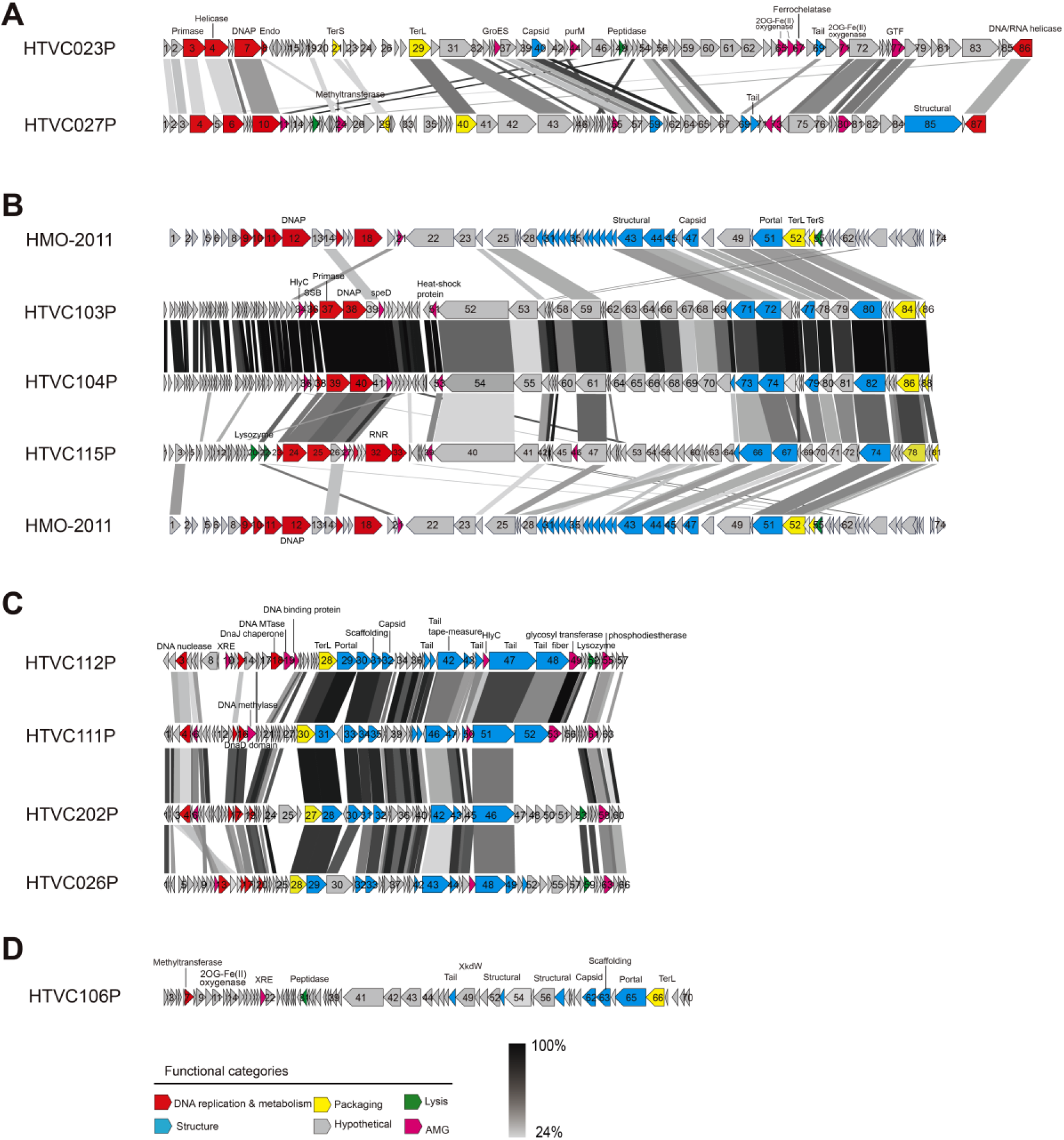
Genomic organization and functional annotation of distinct pelagiphage genera. (*A*) HTVC023P-type pelagiphage genomes, (*B*) HTVC103P-type pelagiphages are compared to SAR116 phage HMO-2011, (*C*) HTVC111P-type pelagiphage genomes, (*D*) HTVC106P genome. Predicted ORFs are indicated by arrows and color-coded according to their putative biological function. Homologous genes were connected by dashed lines. The color of the shading connecting homologous genes indicate the level of amino acid identities between genes. The arrows also designates the direction of transcription. Abbreviation: RNAP, RNA polymerase; SSB, single-stranded DNA binding protein; endo, endonuclease; DNAP, DNAP polymerase; exon,exonuclease; MazG, pyrophosphatase; DNA cytosine methyltransferase; FkbM family methyltransferase; TerS, terminase small subunit; TerL, terminase large subunit; GroEs, Co-Chaperonin GroES; HlyC, toxin-activating lysine-acyltransferase; purM, phosphoribosylaminoimidazole synthetase; GTF, Glycosyltransferase.

ORFs encoding the proteins necessary for phage DNA replication, packaging, morphology and lysis were identified. Although HTVC023P and HTVC027P resemble podoviruses in morphology, only few ORFs in their genomes have the best hit to other phages in the *Podoviridae* family. In the DNA replication region, both HTVC023P-type pelagiphages contain a DNA polymerase gene, a DNA helicase gene and a few function-unknown ORFs that are homologous to genes from siphoviruses, including *Dinoroseobacter* phage vB_DshS-R5C, *Proteobacterial* phage phiJL001 and several *Yuavirus* siphophages, with low amino acid identity (ranging from 26 to 39%). Very few putative structural proteins could be identified with weak homology to other phage structural proteins. For example, HTVC027P ORF85 is homologous to the putative structure protein from *Cellulophaga* phage 18:3, and two others ORFs (HTVC023P ORF69 and HTVC027P ORF69) have small regions of homology to the putative tail fiber genes from some phage genomes. These results imply the existence of a novel set of phage structural proteins in the HTVC023P and HTVC027P genomes. Terminase large and small subunit (TerL and TerS) genes involved in DNA packaging were predicted, which are more closely related to TerL genes in some bacterial genomes. Both HTVC023P and HTVC027P harbor a GroES gene that encode a 10 kDa co-chaperonin. Many bacteria contain the GroEL/GroES molecular chaperonin system that is responsible for proper folding of many proteins, thus playing an important role in cell growth and cellular phage assembly (23, 24). Bacteriophage encoded cochaperonins were identified and studied in some phage genomes (25, 26). Both HTVC023P and HTVC027P GroES genes do not show significant homology to any phage GroES, while being mostly related to GroES sequences retrieved from marine viromes (27). Sequence analyses suggests that GroES in HTVC023P and HTVC027P are clustered with GroES clusters 14 and cluster 1, respectively, which were among the most abundant GroES clusters identified from viromic datasets (27).

The result of the DNA polymerase gene phylogeny analysis shows that HTVC023P and HTVC027P DNA polymerases are placed with clade III DNAP genes identified in a previous shotgun metaviromes study (28), and DNA polymerases from vB_DshS-R5C, *Yuavirus* siphoviruses and phiJL001 are more distantly related (Fig. 1*C*). Clade III DNA polymerases accounted for 77% of all identified DNA polymerases from the Chesapeake Bay, Gulf of Maine and Dry Tortugas (28). It was previously speculated that Clade III DNA polymerases are likely from lysogenic phages (28), whereas our study shows that this DNA polymerase group is related to lytic podoviruses represented by HTVC023P-type pelagiphages.

Gene-content-based network analysis reveals that 443 viral sequences (≥20kb) from diverse ocean regions were grouped into a viral cluster (VC_009) with HTVC023P-type pelagiphages (Fig. 3), suggesting that that close relatives of HTVC023P-type pelagiphages exhibit globally distribution pattern. Phylogenic analysis based on the VC_009 DNA polymerase sequences reveals a high level of diversity (Fig. 4). We notice that GOV populations grouped with HTVC023P-type pelagiphages are exclusively from GOV viral clusters VC_6 and VC_8 (5). VC_6 and VC_8 were two of the most globally abundant viral clusters identified in GOV study (5). Genomic analysis reveals the genetic relatedness and genome synteny between HTVC023P-type pelagiphages and representative contigs from GOV VC_6 and VC_8, showing a high degree of synteny (Fig. 5*A*). Approximately half of the contigs in GOV VC_6 and VC_8 share more than 40% genes with HTVC023P-type pelagiphages and most of the remaining contigs share more than 20% genes with HTVC023P-type pelagiphages. In most cases, the low percentage of shared genes between viral populations and HTVC023P-type pelagiphages is due to some environmental viral sequences covering the nonconserved variable phage genome regions (data not shown). These results suggest that HTVC023P-type pelagiphages and most viral populations from GOV VC_6 and VC_8 can be grouped at the genus/subfamily-level. The phylogenetic analysis reveals that all GOV VC_6 and VC_8 DNA polymerases are clustered with HTVC023P-type DNA polymerases, with 36% to 87% amino acid identity (*S1 Appendix*, Fig. S1).

**Fig. 3.**
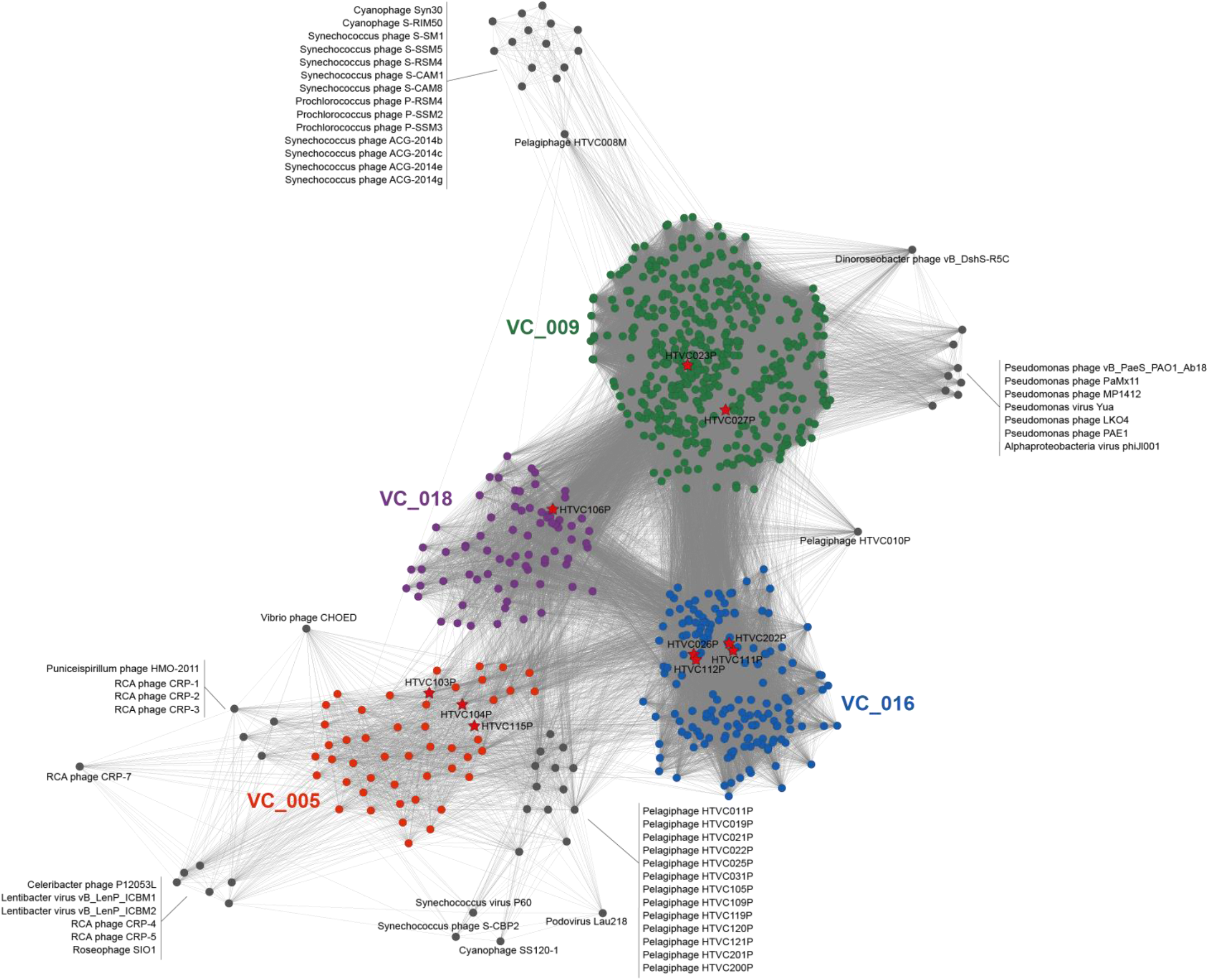
Gene-content-based viral network of pelagiphages, related bacteriophages from NCBI, and related environmental viral sequences from Mediterranean DCM (MedDCM) fosmids, GOV and GOV2.0. The nodes represent the viral genomic sequences. The edges represent the similarities score between genomes based on shared gene content. Viral genomes that belong to different viral clusters are indicated by different colors. For clarity, only environmental viral sequences grouped with ten pelagiphages were presented, and only bacteriophages have genome-genome similarity score of ≥1 with these four phage clusters were presented. Viral clusters generated by vConTACT2 are provided in Dataset S2 in the supplemental material.

**Fig. 4.**
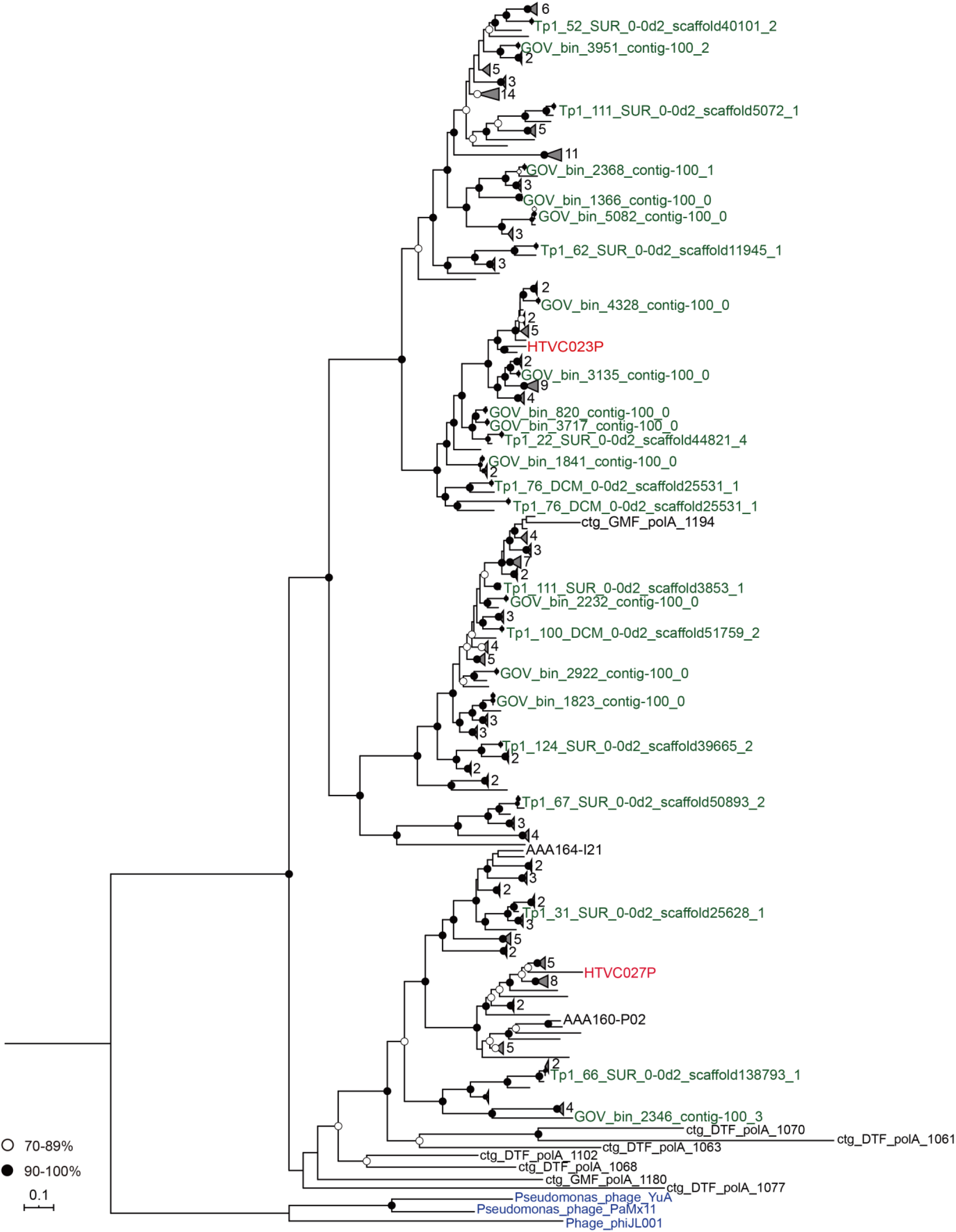
Maximum-likelihood tree of DNA polymerases from the viral cluster VC_009 generated by vConTACT v.2.0 in this study. HTVC023P-type pelagiphages, GOV populations (VC_6 and VC_8) and outgroups are indicated in green, red, and blue, respectively. The names of the GOV2.0 populations are omitted in the tree.

**Fig. 5.**
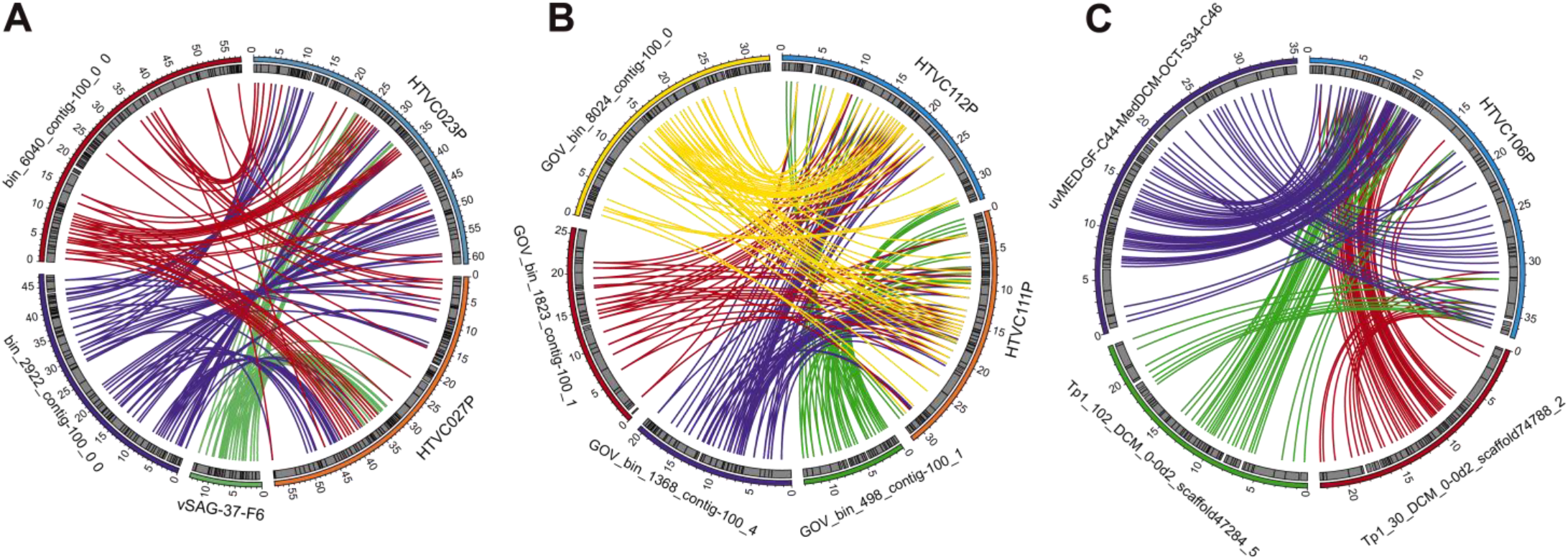
Circos comparison plot indicating the genome comparison between pelagiphages and environmental viral fragments. (*A*) Comparison of HTVC023P-type genomes, vSAG 37-F6, and representative viral populations in VC_6 and VC_8. (*B*) Comparison of HTVC111P-type genomes and representative viral populations in VC_41. (*C*) Comparison of HTVC106P genome and representative viral populations in VC_67. Each coloured segment represents a phage genome or viral fragment with the numbers on the external surface indicating genome size in kb. Homologous genes shared between genomes are connected by color lines. Only the relatedness between pelagiphages and other sequences is indicated.

These two pelagiphages also show high homology with a viral single-amplified genome contig, vSAG 37-F6 (13) (Over 80% of the predicted proteins in vSAG 37-F6) (Fig. 5*A*). The vSAG 37-F6 population was reported to be closely related to GOV VC_6 and VC_8 and has been shown to be abundant in several oceanic regions (13). SAR11 was recently predicted as putative host of vSAG 37-F6 population by single-cell genomics (29). However, before our study, phage-host system of these extremely important viral clusters still remained unavailable. It is noteworthy that in an earlier study, homologs of HTVC023P-type genomes were also found in single cell genomic analyses of *Verrucomicrobia* and *Bacteroidetes* (AAA160P02 and AAA164-I21) (*S1 Appendix*, Fig. S2) (12), suggesting that members of the HTVC023P-type group may infect different taxonomic groups of bacteria. Further investigation based on culture-independent or culture-dependent studies are required to explore the diversity and infected hosts of this important viral group.

Taken together, close relatives of HTVC023P-type pelagiphages have been previously identified from some culture-independent studies and were revealed to be extremely abundant. These two HTVC023P-type pelagiphages are first known cultured representatives of this important viral group.

### Homology to the HMO-2011-type phage group

Three pelagiphages, HTVC103P, HTVC104P and HTVC115P are closely related, belonging to a novel HTVC103P-type group (Figs. 1*B* and 2*B*). Approximately 15% of their ORFs were assigned with putative functions. HTVC103P-type pelagiphages exhibit novel genomic architectures, containing two functional modules, including a DNA replication module and a phage structural and packaging module (Fig. 2*B*). In the DNA replication module, DNA polymerase, DNA primase and single-strand binding protein were predicted from all three HTVC103P-type pelagiphage genomes. The closest homologs of HTVC103P-type DNA polymerases and primases from isolated phages are those found in members of the *Autographivirinae* subfamily and Cobavirus group roseophages. Interestingly, HTVC103P and HTVC104P both share 12 genes with SAR116 phage HMO-2011 and HTVC115P shares 16 genes with HMO-2011. Most of the HMO-2011 homologs are in the structural and packaging modules, including genes encoding capsid, portal and terminase (Fig. 2*B*). In addition, there is considerable amino acid identity and conserved gene synteny between HTVC103P-type pelagiphages and HMO-2011 in this region (27-68% amino acid identity). Gene content-based network analysis also reveals the relatedness between HTVC103P-type pelagiphages and HMO-2011-type phages (Fig. 3). These results suggest that HTVC103P-type genomes are probably composed of a DNA replication module and a phage structural and packaging module with distinct evolutionary origins and histories. These results indicate that horizontal gene exchange of the function module among phages may play an important role in driving evolution and genetic diversity of pelagiphages. It is likely that the transfer of a set of structural or DNA replication machinery genes occurred when divergent phages infected the same host cell or when there was contact between a resident prophage and an invading phage. Further investigation are required to illuminate the evolutionary trajectories of this novel phage group.

The DNA polymerase gene based phylogeny reveals that HTVC103P-type DNA polymerases are grouped with clade I DNA polymerases and are more distantly related to DNA polymerases from other *Autographivirinae* phages and Cobavirus roseophages (Fig. 1*C*). Clade I was another abundant DNA polymerase clade that was previously identified from shotgun metaviromes (28). In contrast, the structural genes of HTVC103P-type pelagiphages are most similar to those in HMO-2011-type phages. This result suggests that a phylogenetic approach based on a single gene has a limitation in revealing the evolutionary relationship among various phages. A gene content-based network can be used as a complement. Network analysis shows that a group of environmental viral fragments (49 sequences, ≥20kb) were clustered with HTVC103P-type pelagiphages, forming a viral cluster VC_005 (Fig. 3). VC_005 shows distant relatedness to the HMO-2011-type group (Fig. 3). The DNA polymerase gene phylogeny reveals that the DNA polymerase genes of VC_005 all cluster with HTVC103P-type pelagiphages and are distinct from HMO-2011-type DNA polymerases (*S1 Appendix*, Fig. S3).

### The new HTVC111P-type group and HTVC106P pelagiphage

The HTVC111P-type phage group currently comprises pelagiphage HTVC111P, HTVC112P, HTVC026P and HTVC202P. Within this group, 45% to 67% of genes were shared (Fig. 1*B*). Genes responsible for phage replication, morphology, packaging, and lysis were identified from HTVC111P-type pelagiphage genomes with homology to genes from diverse bacteria and bacteriophages (Fig. 2*C*). The DNA polymerase gene was not found in the HTVC111P-type pelagiphage genomes and structural genes show very weak similarity to proteins from other known phage genomes, suggesting that this group of phages contain novel morphogenesis modules and may rely more on host replication system. The TerL gene in the HTVC111P-type genomes also show no significant similarity to other phage TerL genes.

Pelagiphage HTVC106P also lacks a clear relationship to any known phage isolates. Of 70 predicted ORFs in HTVC106P, approximately 40% have homologs in other organisms in the NCBI-RefSeq database and only 11 ORFs were assigned with putative functions (Fig. 2*D*). For the remaining ORFs, most were highly similar to genes found only in metagenomic sequences. The ORFs assigned with functions are involved in DNA processing, virion morphogenesis, DNA packaging and lysis. The DNA replication genes were not identified in the HTVC106P genome, suggesting that HTVC106P may rely more on the host replication system or contain a novel replication module. The morphogenesis genes of HTVC106P show homology to some phages; for example, the HTVC106P portal protein, capsid and scaffolding protein exhibit distant homology with those of *Bruynoghevirus* phages, and the HTVC106P tail fiber protein and some other structural proteins are homologous with those in pelagiphage HTVC010P. HTVC106P contains a TerL related to the TerL in pelagiphage HTVC010P (49% amino acid identity), suggesting that HTVC106P possibly shares a conserved DNA packaging strategy with HTVC010P.

The close relatedness between HTVC111P-type pelagiphages, HTVC106P and some metagenomic viral fragments are also revealed by network analysis, with 165 and 75 sequences (≥20kb) grouped with HTVC111P-type pelagiphages and HTVC106P, respectively (see VC_016 and VC_018 in Fig. 3). Phylogeny of capsid protein sequences reveals a high level of diversity of these two viral clusters (*S1 Appendix*, Figs. S4 and S5). Network analysis also reveals the affiliation of pelagiphages with previouly identified viral clusters in GOV. The GOV viral populations grouped with HTVC111P-type pelagiphages and HTVC106P are exclusively from GOV viral cluster VC_41 and VC_67, respectively (5). Approximately 80% of the viral populations (≥20kb) of VC_41 and 70% of the viral populations (≥20kb) of VC_67 are clustered with HTVC111P-type pelagiphages and HTVC106P, respectively. Genomic analysis reveals the genetic relatedness and genome synteny between HTVC111P-type pelagiphages and representative contigs from GOV VC_41, HTVC106P and representative contigs from VC_67 (Fig. 5*B*, 5*C*). The hosts of GOV VC_41 and VC_67 were not predicted yet (5). Most viral populations in GOV VC_41 and VC_67 share more than 20% genes with HTVC111P-type phages and HTVC106P, respectively, indicating a possible relationship between these phages at the subfamily-level.

### Pelagiphage lytic and lysogenic developmental strategies

Although the integrase gene was not identified from all ten pelagiphages (described above), a search for integrase genes reveals that a total of 27 environmental viral sequences closely related to HTVC106P were found to contain a tyrosine integrase gene (PFAM, PF00589) (*S1 Appendix*, Fig. S6). Phage integrase mediates the site-specific recombination between phage sequence and bacterial sequences(30, 31). In addition, most of these integrase-containing viral sequences possess a sequence identical to SAR11 tRNA sequences (tRNA-Thr or tRNA-Asn) (*S1 Appendix*, Fig. S6), which are likely to be the phage integration sites. In contrast, no identifiable integrases were found from environmental viral sequences closely related to other three phage groups. Although no intact prophages have been found in SAR11 genomes, lysogenic infection has been reported in HTVC019P-type pelagiphages (22). Our analysis suggests that a portion of the HTVC106P-type pelagiphages can infect the hosts via the lysogenic cycle, while other three pelagiphage types may have a strict lytic life strategy. These findings indicate that pelagiphage have diverse life strategies and lytic infection strategy is presumably the predominant form of pelagiphage-SAR11 interaction.

### Pelagiphages dominate the marine viromes

The ecological importance of pelagiphages are reflected by their sheer abundance and ubiquity, as well as the predominance of their hosts in the ocean. In recent years, the marine viromic reads increased exponentially, providing valuable resources for assessing the relative abundance and distribution pattern of important viral groups in the ocean (5, 6, 32). In this study, a total of 174 marine virome datasets from various oceanic sites were downloaded for the viromic fragment recruitment analysis (*S1 Appendix*, Table S1). We mainly used the reciprocal best-hit strategy to estimate the relative abundance of phage groups. Instead of accessing the relative abundance of each viral genome by recruiting viromic reads with a high nucleotide identity (>95% or 90%) (9, 10, 13, 33, 34), reciprocal recruitment method estimates the relative abundance of different phage groups at the genus or subfamily level. Due to the fact that a viral taxonomic group comprise evolutionarily diverse genotypes, reciprocal recruitment analysis can obtain an estimation of the relative abundance of important phage groups.

It is striking that within the top ten most abundant known phage groups at each marine viromic dataset, approximately half were represented by pelagiphages (Fig. 6 and Dataset S1), suggesting that pelagiphages are abundant components of the marine viral communities. The HMO-2011-type group, some cyanophage- and roseophage-represented groups were also abundant throughout the world’s ocean (Fig. 6). The new HTVC023P-type group was the most abundant viral group in the majority (80%) of the analyzed viromic datasets, exceeding the HMO-2011-type group and the HTVC010P-type group (Fig. 7). On average, the HTVC023P-type group is 2.8 and 2.2 times more abundant than the HMO-2011-type group and the HTVC010P-type group, respectively. The reads assigned to the HTVC023P-type group accounted for 0.55% to 7.59% of total viromic reads in various oceanic viromes, suggesting that the HTVC023P-type phage group dominates the marine viromes (Dataset S1).

**Fig. 6.**
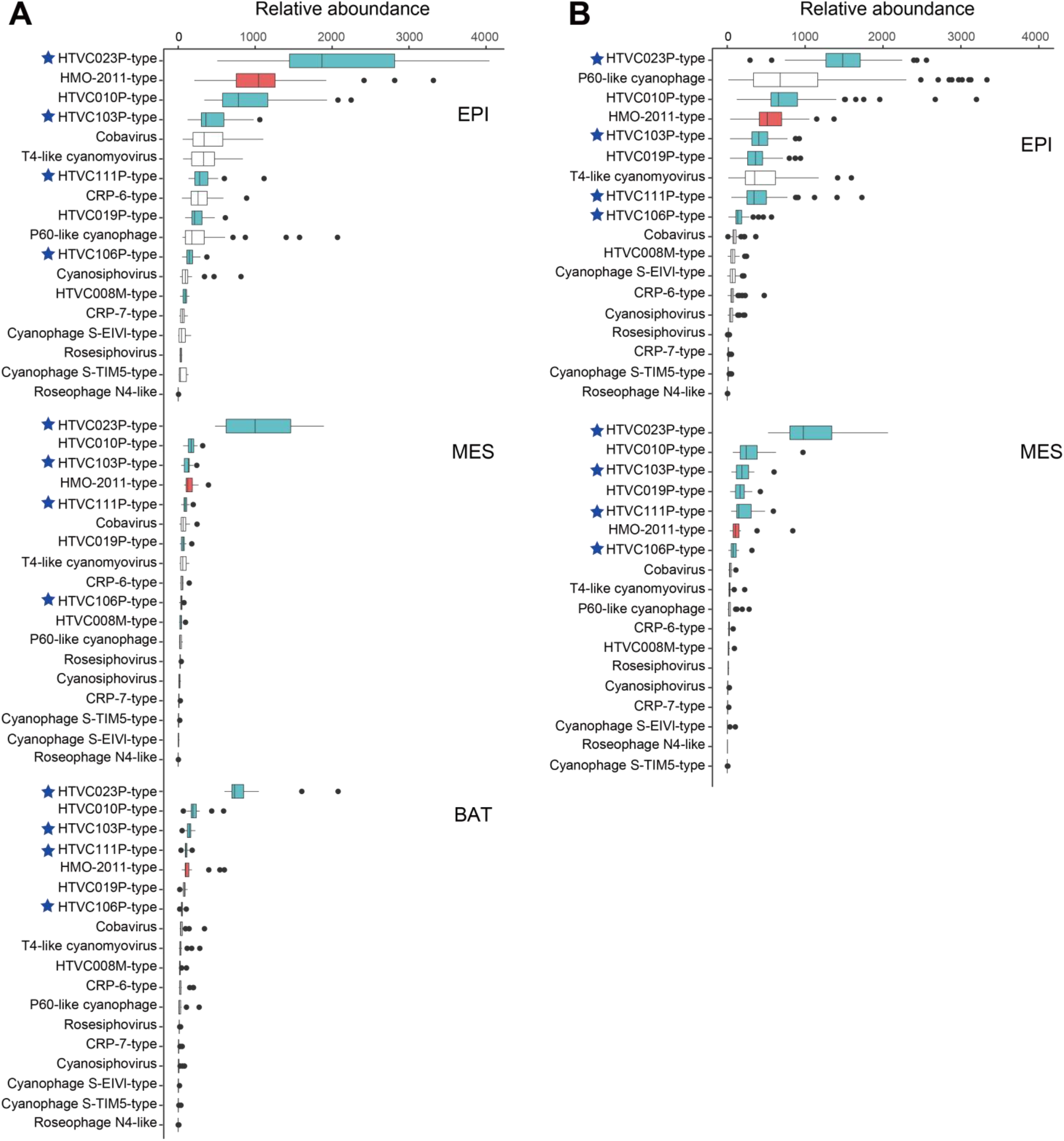
Box plots indicate the relative abundance of major phage groups in different marine viromic datasets. Normalized read recruitment is depicted as the number of reads recruited per kilobase of the genome per billions reads in the dataset. Pelagiphage represented groups are colored in blue; the HMO-2011-type group is colored in red. Pelagiphage groups identified in this study are marked with blue stars. (*A*) Relative abundance of major phage groups in epipelagic, mesopelagic and bathypelagic samples in POV, MES, SPV and IOV. (*B*) Relative abundance of major phage groups in epipelagic, mesopelagic and bathypelagic samples in the Global Oceans Viromes (GOV). EPI, Epipelagic; MES, Mesopelagic; BAT, Bathypelagic.

**Fig. 7.**
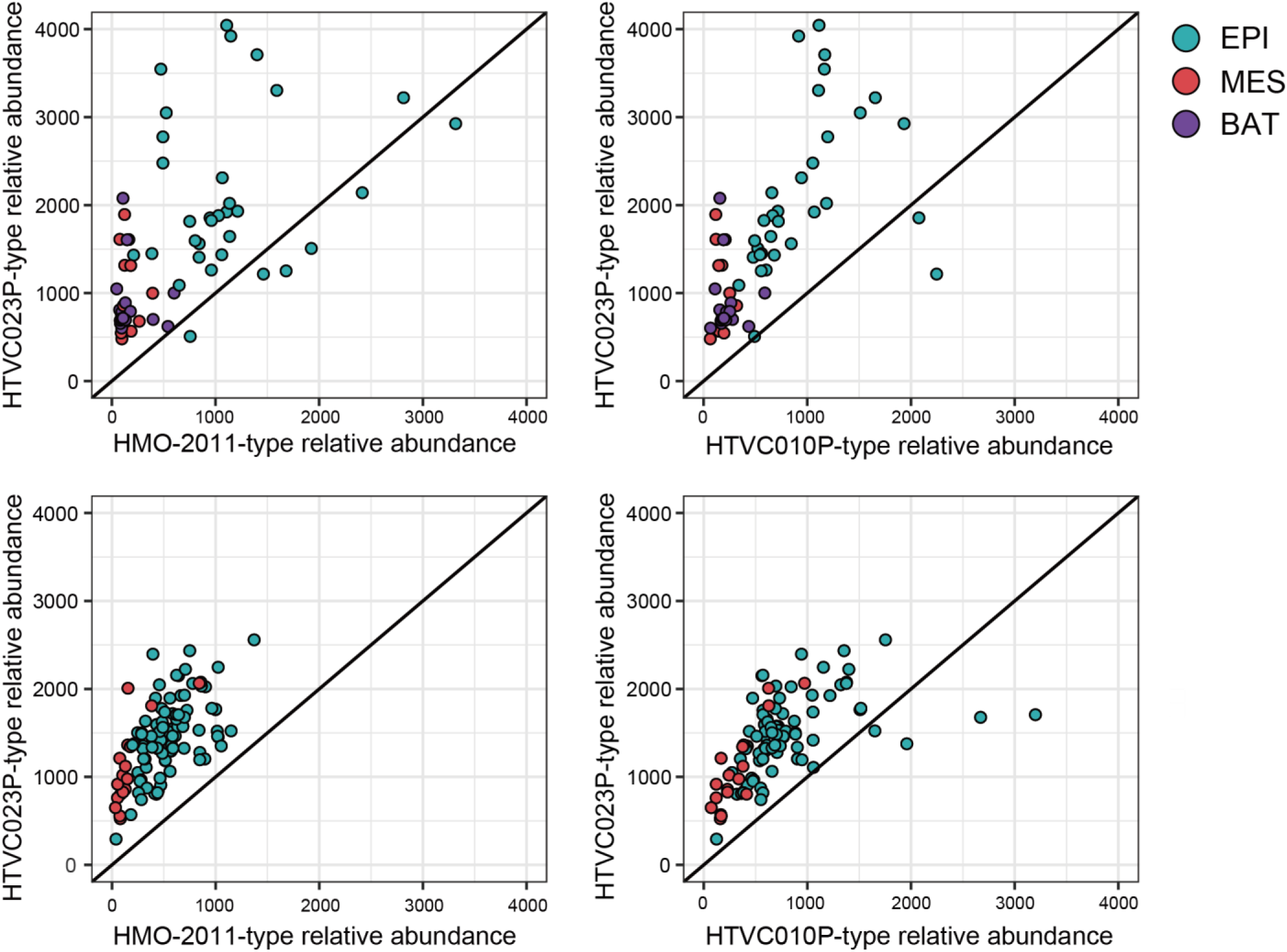
Comparison of the relative abundance between HTVC023P-type and HMO-2011-type, HTVC023P-type and HTVC010P-type. Upper panel: comparison of the relative abundance in POV, MES, SPV and IOV. Lower panel: comparison of the relative abundance in GOV. EPI, Epipelagic; MES, Mesopelagic; BAT, Bathypelagic.

The HTVC103P-type group ranked among the five most abundant viral groups in most viromic datasets. A remarkable feature of HTVC103P-type pelagiphages is that they harbor a set of structural genes that are homologous to those in HMO-2011-type phages. HMO-2011-type phages were highly represented in some ocean viromes, and the known host of HMO-2011-type phages currently comprise SAR116 and RCA roseobacters (15, 35). The structural gene homology between HTVC103P-type pelagiphages and HMO-2011-type phages raise the possibility that a portion of the viromic reads that were previously assigned to the HMO-2011-type group might be assigned to the HTVC103P-type group when HTVC103P-type genomes are included in the analysis. It was then estimated that the reads that were assigned to the HMO-2011-type group decreased by approximately 12% when HTVC103P-type pelagiphage genomes were included in the analysis (data not shown), thus demonstrating that the HTVC103P-type group contributed to the abundance of HMO-2011-type phages in previous studies. When more related phage isolates are available, the analysis on phage relative abundance might be different to some extent.

Among all known pelagiphage groups, the HTVC112P-type and HTVC106P-type were less abundant than other pelagiphage groups, but they were still abundant and globally distributed. In the upper waters, the HTVC111P-type appeared to be as abundant as the HTVC019P-type group and the T4-like cyanomyoviruse group.

The above results further support that pelagiphages are extremely abundant and widely distributed in the ocean. Considering the ubiquity and dominance of SAR11 bacteria in the ocean, they are prone to be attacked by viruses. The vast population sizes of SAR11 support the thriving of pelagiphages. The prevalence of pelagiphages in marine viromes suggests that SAR11 populations are under intense phage infection pressure. Podoviruses act as a primary contributor to SAR11 mortality and exert major control on the abundance and dynamics of SAR11 population. Although there is presumably significant cell loss of SAR11 populations due to the viral predation, as described by the King of the Mountain (KoM) hypothesis, the high recombination frequencies of SAR11 may also influence the distribution of phage-defense alleles, maintaining the coexistence of a high abundance of host and phages (16).

Among all known pelagiphage represented phage groups, the dominance of HTVC023P-type phages raises the question of what characteristics make them most successful. It presumably links to their biological traits; that is, they are likely to have higher infection efficiency (faster replicating) or they are more competitive when competing with other types of phages for hosts in complex viral assemblages.

### Vertical profiles of pelagiphages

We compared the relative abundance of major marine phage groups at different water depths. There were significant variations in the relative abundance of most phage groups on the vertical scale (Fig. 6*A*, *B*). The relative abundance of the HTVC023P-type group appeared lower in mesopelagic (200 to 1000 m) and bathypelagic (1000 to 4000 m) samples than in epipelagic samples. Considering the cellular contamination reported in the bathypelagic viromes (5), it is possible that the HTVC023P-type exhibited comparable abundance throughout the water column. Remarkably, we observed that the HTVC023P-type group far exceeded other phage groups in the deep waters (200 to 4000 m) (Figs. 6 and 7). For example, in bathypelagic viromes, the HTVC023P-type group was on average 8 and 6 times more abundant than the HMO-2011-type and HTVC010P-type, respectively. Moreover, four of the five most abundant viral groups in mesopelagic and bathypelagic samples were represented by pelagiphages. In congruence, SAR11 is abundant throughout the oceanic water column and is abundant in coastal and open ocean (18, 36, 37); In deep waters, SAR11subclades Ic, IIb, and Vb dominate the SAR11 populations (38, 39); thus, in the deep ocean, members of these phage groups are likely to infect these deep ocean SAR11 ecotypes. These results suggest that pelagiphages could be as important in deep ocean ecosystems as they are in the upper ocean. In contrast, the HMO-2011-type group only predominates the upper ocean, which is consistent with previous study (40). HMO-2011-type phages were found infecting SAR116 and RCA rosephages, which mainly occupy the upper ocean (15, 35). Furthermore, we observed that the relative abundance of all pelagiphage groups did not exhibit significant variation between the coastal viromes and noncoastal viromes (*S1 Appendix*, Fig. S7).

### Conclusions

The 10 new pelagiphages that were described in this study greatly expand our current knowledge on the abundance, diversity, distribution and genomic evolution of viruses that infect SAR11 bacteria and further reinforce their ecological importance. Metagenomic recruitment analyses demonstrate that all these pelagiphage represented phage groups exhibit global distribution pattern and the HTVC023P-type group is the most dominant known viral group in the ocean. The predominance of these pelagiphages in marine viromes suggests that they could play an important role in controlling SAR11 population dynamics and influencing global carbon cycling. These new pelagiphages and their hosts will serve as useful model systems subjected to the further investigations of various interactions between pelagiphages and their hosts, phage driven host evolution and dynamics, as well as the potential ecological impact of pelagiphages. It will also be interesting to study the mechanisms explaining the dominance of major phage groups. This study is another example of how phage isolation can improve the interpretation of marine viromic datasets, thus highlighting the irreplaceable power of culture-dependent phage isolation and cultivation in the study of marine virus functions and diversity. So far, all current known pelagiphage isolates were isolated from strains from the SAR11 Ia subclade. Future isolation of pelaiphages that infect other SAR11 subclades may reveal more novel phage lineages.

## Methods

### SAR11 strains, media, and growth conditions

SAR11 strains *Pelagibacter* HTCC7211 and *Pelagibacter* HTCC1062 were grown in an artificial seawater-based mediun amended with 1 mM NH_4_Cl, 100 μM KH_2_PO_4_, 1 μM FeCl_3_, 100 μM pyruvate, 50 μM glycine, 50 μM methionine and excess vitamins (21). HTCC7211 and HTCC1062 were grown at 17 ℃ and 20 ℃, respectively. *Pelagibacter* FZCC0015 was isolated from Pingtan coast in 2017, detailed information on FZCC0015 has been described in earlier work (22). FZCC0015 was grown in a natural seawater-based medium amended with 100 μM pyruvate, 50 μM glycine, 50 μM methionine and excess vitamins at 23 ℃.

### Source waters and pelagiphage isolation

Water samples were collected from a variety of oceanic sampling stations (Table 1). To obtain the cell-free fraction, the samples were filtrated through 0.1 μm-pore-size filters. The filtrates were stored in the dark at 4 °C until used for phage isolation. Isolation procedures for pelagiphages were described in detail previously (16, 22). Briefly, 0.1μm filtered samples were inoculated with SAR11 cultures. Cell growth was monitored using a Guava EasyCyte cell counter (Millipore, Guava Technologies). When a decrease in cell densities was detected, the presence of phage particles was confirmed by epifluorescence microscopy. Purified pelagiphage clones were obtained by using the dilution-to-extinction method (35). The purity of pelagiphages was verified by whole-genome sequencing.

### Morphological analysis by transmission electron microscopy

Representative pelagiphages were observed by transmission electron microscopy (TEM). Pelagiphage lysates were filtered through 0.1μm pore-size filters and then concentrated using Amicon Ultra Centrifugal Filters (30-kDa, Merck Millipore). Concentrated phage particles were absorbed onto copper grids in the dark, negatively stained with 2% (wt/vol) uranyl acetate for two minutes, and air-dried. Samples were observed using a Hitachi transmission electron microscope at an acceleration voltage of 80 kV.

### Phage DNA preparation, genome sequencing and functional annotation

Phage lysate preparation and concentration were carried out as described in Zhang and colleagues(35). Briefly, 250 ml of each phage lysate was filtered through 0.1 μm Supor membrane to remove cells and cell debris. Phage lysates were concentrated by centrifugal filtration using Amicon Ultra-15 30-kDa filters (Merck Millipore, Cork, Ireland). Phage genomic DNA was prepared using a formamide, phenol/chloroform extraction protocol(41) and sequenced on an Illumina paired-end HiSeq 2500 platform. The raw reads were quality-filtered, trimmed and de novo assembled with default settings using CLC Genomic Workbench 11.0.1 (QIAGEN, Hilden, Germany). The remaining gaps in each pelagiphage genome were closed by Sanger sequencing of PCR products.

Prodigal(42) and GeneMark (43) were used for phage open reading frames (ORFs) prediction. The translated open reading frames (ORFs) were used as BLASTP queries to search against the NCBI nonredundant (nr) and NCBI-Refseq database. Putative functions were assigned to ORFs based on their homology to proteins of known function. In this study, genes with ≥25% amino acid identity, ≥ 50% alignment coverage of the shortest protein, and an E-value cutoff ≤1E-3 were considered to be putative homologues. A PFAM database search was performed to identify conserved protein domains. HHPred was also employed to identify the distant protein homologs. tRNA prediction was performed using the tRNAscan-SE program (44). Comparative genome map and connections between homologous genes were visualized using CIRCOs(45).

The genomic sequences of the ten pelagiphages have been deposited in GenBank under the accession numbers MN698239 to MN698248.

### Network analysis

Protein sequences from a total of 2591 bacterial viruses’ genomes were downloaded from NCBI-RefSeq (v96). 927 viral genomic sequences (≥20kb, ≥20% shared gene with any pelagiphage genome) from metagenomic fosmids, GOV and GOV2.0 datasets were also included in the network analysis(5, 6, 9). All proteins were compared using all-verses-all BLASTP (e-value ≤1E-5, bitscore ≥50). Protein clusters (PCs) were defined using the Markov clustering algorithm (MCL) (46). vConTact 2.0 was then used to calculate a similarity score between every pair of genomes based on the number of PCs shared between two sequences and all pairs using the hypergeometric similarity (47). The network was created using Cytoscape v.3.5.1 (48).

### Phylogenetic analysis

A phylogenetic tree of DNA polymerase family A domain sequences was constructed to evaluate the evolutionary relationship among pelagiphages and other diverse phages. Alignment for the DNA polymerase family A domain was constructed with MUSCLE (49) and edited with Gblock (50). The alignment was evaluated for optimal amino acid substitution models using ProtTest (51), and run with RAxML v8 (52) with a bootstrap of 500.

In order to evaluate the phylogenetic relationship among pelagiphages and environmental viral sequences, we constructed maximum likelihood phylogenetic trees of DNA polymerases and capsid proteins. Sequence alignments and editing were performed using MUSCLE (49) and Gblocks (50), respectively. Maximum-likelihood phylogenetic trees were constructed using FastTree 2.1 (53) with WAG substitution model for amino acids.

### Search for integrase genes in pelagiphage related viral sequences

Environmental viral genomic sequences from the viral clusters VC_009, VC_005, VC_016 and VC_018 that generated by vConTact 2.0 in this study were used for analysis. To identify integrase genes, hmmbuild was used to build HMM files from phage integrase and recombinase domains. The program hmmsearch was then used to identify the putative integrase genes by searching HMM files against the pelagiphage related environmental viral sequences. Viral sequences containing the putative integrase were subjected to manual inspection and comparative genomic analyse. Putative integration sites were identified by searching against known SAR11 genome sequences using BLASTn.

### Metagenomic fragment recruitment analyses

Marine viromic datasets that were used for accessing phage relative abundance include Pacific Ocean Virome (POV), Scripps Pier Virome (SPV), India Ocean Virome (IOV), Malaspina Expedition virome (ME) and Global Oceans Viromes (GOV) (*S1 Appendix*, Table S1). The fragment reciprocal recruitment method was described in detail in a previous study (35). The analysis procedure is summarized as follows:

1. Each of themarine viromic reads was searched as a query against the NCBI-RefSeq viral database (release 88), 18 recently published HTVC019P-type pelagiphage and RCA phage genomes (22, 35), 6 newly sequenced HTVC010P-type pelagiphage genomes and 10 pelagiphage genomes reported in this study using DIAMOND BLASTx (e-value cutoff of ≤1E-3, bitscore cutoff ≥40). Reads with homology to viral sequences were retained for the subsequent analysis.
2. The reads were assigned to the best-hit virus or best-hit bacteria using BLASTx against RefSeq viral database, RefSeq bacterial database, 11 HTVC019P-type pelagiphages, 7 RCA phage genomes, 6 new HTVC010P-type pelagiphages and 10 newly sequenced pelagiphages.
3. Reads that returned a best-hit of the query genome from the bacteriophage genomes included in the relative abundance analyses (*S1 Appendix*, Table S2) were identified and extracted from the viromic datasets.
4. The relative abundances of each phage group were calculated and normalized as the number of reads recruited to the group normalized to the total number of base pairs in the virome and the average genome size (Reads mapped per kb per millions of reads, RPKM).

Due to the large amount of sequencing data in the Global Oceans Viromes (GOV) datasets (>500G), a different metagenomic analysis strategy was used to determine the relative abundances of different phage groups in the GOV. GOV reads were recruited onto the phage genomes (*S1 Appendix*, Table S2) using BLASTx with an e-value cutoff of 1E-10. If a read was recruited to more than one phage genome, the read was associated with the phage that provided the highest bitscore. RPKM was also used to calculate and normalize the relative abundances of each phage group in each virome datasets in GOV.

### Data Availability Statement

The genomic sequences of the ten pelagiphages have been deposited in GenBank under the accession numbers MN698239 to MN698248. The data that support the findings of this study are available upon request from the corresponding authors.

## Acknowledgments

The work has been supported by NSFC grant 41706173. We thank Sijun Huang for providing the water samples. We thank Chen Li and Sun Jing for their assistance in TEM.

